# EPIKOL, a chromatin-focused CRISPR/Cas9-based screening platform, to identify cancer-specific epigenetic vulnerabilities

**DOI:** 10.1101/2021.05.14.444239

**Authors:** Ozlem Yedier-Bayram, Bengul Gokbayrak, Ali Cenk Aksu, Ayse Derya Cavga, Alisan Kayabolen, Ezgi Yagmur Kala, Goktug Karabiyik, Rauf Günsay, Tunc Morova, Fırat Uyulur, Nathan A. Lack, Tamer T. Önder, Tugba Bagci-Onder

## Abstract

Dysregulation of the epigenome due to alterations in chromatin modifier proteins commonly contribute to malignant transformation. To discover new drug targets for more targeted and personalized therapies, functional interrogation of epigenetic modifiers is essential. We therefore generated an epigenome-wide CRISPR-Cas9 knock-out library (EPIKOL) that targets a wide-range of epigenetic modifiers and their cofactors. We conducted eight screens in two different cancer types and showed that EPIKOL performs with high efficiency in terms of sgRNA distribution, depletion of essential genes and steady behaviors of non-targeting sgRNAs. From this, we discovered novel epigenetic modifiers besides previously known ones that regulate triple-negative breast cancer and prostate cancer cell fitness. With further validation assays, we confirmed the growth-regulatory function of individual candidates, including SS18L2 and members of the NSL complex (KANSL2, KANSL3, KAT8) in triple negative breast cancer cells. Overall, we show that EPIKOL, a focused sgRNA library targeting approximately 800 genes, can reveal epigenetic modifiers that are essential for cancer cell fitness and serve as a tool to offer novel anti-cancer targets. With its thoroughly generated epigenome-wide gene list, and the relatively high number of sgRNAs per gene, EPIKOL offers a great advantage to study functional roles of epigenetic modifiers in a wide variety of research applications, such as screens on primary cells, patient-derived xenografts as well as *in vivo* models.

## INTRODUCTION

Epigenetic modifications regulate gene expression and are altered by developmental and environmental cues [1]. Strict epigenetic control is required during embryogenesis, differentiation, cell fate decisions and maintenance of cell identity [2]. Dysregulation of the epigenome has emerged as an important mechanism contributing to various pathologies including tumorigenesis. Epigenome-level alterations pave the way for pre-malignant cells to acquire cancer hallmarks including aggressiveness, environmental adaptation and resistance to therapy [3]. For example, in many cancer types, DNA hypomethylation activates proto-oncogene expression, whereas DNA hypermethylation represses tumor-suppressor expression [4-8]. Similarly, acetylation of histones leads to relaxed chromatin allowing aberrant gene expression whereas methylated histones are often associated with gene silencing [9, 10]. Recent cancer genome sequencing studies revealed mutations in many epigenetic modifiers that are associated with various cancers [11], such as DNMT3A in acute myeloid leukemia [9, 12], IDH1/2 in glioblastoma [13, 14], CREBBP/EP300 in small-cell lung cancer [15] and ARID1A in gastric cancer [16]. These driver mutations are thought to act in part by increasing cellular plasticity during development of malignant tumors. Given this critical role, small molecule inhibitors targeting epigenetic regulators are promising anti-cancer drugs and have shown efficacy in various cancer types [17]. However, first-generation molecules have had limited clinical benefit due to their high toxicity [18, 19]. To overcome these limitations, newer molecules are being developed and tested in clinical trials [20-23]. Using these inhibitors to take advantage of synthetic lethal interactions between epigenetic modifiers, offers a promising therapeutic approach to target the disease in a cancer-specific manner [20, 24-28].

CRISPR/Cas9 technology is a fast, effective and easy-to-use genome engineering method and has drastically accelerated functional genomics research [29]. Its simplicity allows for the generation of multiplexed sgRNA libraries to interrogate gene functions in pooled genome-wide knockout screens [30, 31]. Negative selection screens identified many essential genes in different contexts [32-34] while positive selection screens helped to identify ‘winner’ genes under a given selective pressure [30, 31]. Although genome-wide CRISPR knockout libraries are versatile tools to study various phenotypes simultaneously, the design and execution of such experiments are laborious and expensive. In many cases, secondary screens focusing on the pathways identified in the primary screen are performed to eliminate false-negative results and obtain high confidence leads. Unlike the limited number of sgRNAs per gene in genome-wide libraries, sgRNA numbers per gene can be increased in focused libraries to enhance reliability of the observed phenotype [35]. Therefore, focused sgRNA libraries have emerged as a way to overcome these challenges by reducing the cost and labor and maximizing the yield and signal/noise ratio [36]. Additionally, focused libraries may be advantageous in experimental systems that require clinically relevant models such as primary cells, patient-derived xenografts [37] or *in vivo* models [38-40]. To date, various focused libraries have been generated targeting microRNAs [41], kinases [42, 43], nuclear proteins [37], epigenetic modifiers [44-46] or genes belonging to a certain pathway such as DNA-damage response [47].

Here, we present our focused <u>Epi</u>genetic <u>K</u>nock-<u>o</u>ut <u>L</u>ibrary (EPIKOL), which targets a broader range of epigenetic modifiers and consists of more sgRNAs for each gene when compared to previously published libraries [44-46]. Utilizing this epigenome wide library in screens of two different cancer types, we revealed novel epigenetic modifiers that regulate cancer cell fitness. Demonstrating the suitability of our library for the identification of epigenetic vulnerabilities of cancer cells, we validated several of these genes in triple-negative breast cancers.

## METHODS

### Library Content of EPIKOL

To generate a customized <u>epi</u>genetic <u>k</u>nock-<u>o</u>ut library (EPIKOL), curated epigenetic modifiers in the EpiFactors database were targeted by sgRNAs [48]. In addition to 719 genes that have roles in chromatin-related pathways, 25 genes from different families (nuclear receptors, ABC transporters, apoptosis or metastasis related proteins) were added to serve as internal controls in specific screen setups. 35 essential genes, such as ribosomal protein encoding genes and 80 non-targeting sgRNAs were also included in the library. 35 essential genes were determined through analysis of publicly available screen data of 60 different cell lines obtained from the Genome CRISPR database [49]. Among them, genes that have the highest log2fc were included in the library. Each gene in EPIKOL is targeted by 10 sgRNAs that were chosen from previously established genome-wide CRISPR knock-out libraries [50, 51]. Additional sgRNAs were designed by using CCTop and E-CRISP tools in cases where the total number of sgRNAs did not reach 10 per gene due to overlapping sequences in existing libraries [52]. Genes and sequences of sgRNAs of EPIKOL are available in **Supplementary Table 1**.

### Cloning of EPIKOL

All sgRNAs were synthesized as pooled oligonucleotides (LC Biosciences). Lyophilized oligonucleotides were resuspended and amplified by PCR with following conditions: For 50 µl of total PCR mix, 10 µl of 5X HF buffer (NEB,USA), 1 µl of dNTP mix (10 mM) (Thermo Fisher,USA), 2.5 µl of forward primer (10 µM), 2.5 µl of reverse primer (10 µM), 0.1 µl of resuspended oligomix, 0.5 µl of Phusion High-fidelity DNA polymerase (NEB,USA), 33.4 µl of dH_2_O were added. Thermal cycler conditions were: 30 sec at 98°C for initial denaturation, followed by 20 cycles of (10 sec at 98°C, 20 sec at 63°C, 15 sec at 72°C), 3 min at 72°C for final extension. Two tubes of 50 µl reaction were run on Agarose gel and correct sized bands were gel-extracted. 10 µg of lentiviral backbones lentiCRISPRv2 (Addgene #52961) and lentiGuide-Puro (Addgene #52963) were digested with BsmBI at 55°C for 6 h followed by agarose gel extraction. PCR-amplified oligos and gel-extracted vector backbones were purified with AMPure XP magnetic beads according to manufacturer’s instructions. For ligation reaction, 100 ng of purified backbone, 15-20 ng of purified oligomix, 5 µl of Gibson assembly mastermix (NEB) and dH_2_O up to 10 µl were mixed and incubated at 50°C for 1 h. 1 µl from ligation reaction was added onto 25 µl of electrocompetent cells (Lucigen) and cells were transformed by using Electroporator (Bio-Rad MicroPulser) according to manufacturer’s instructions. 1 ml of recovery medium was added on cells immediately after pulse and incubated at 32°C for 1 h. 10 µl of bacteria culture were taken to 90 µl of LB and serial dilution was performed to estimate library coverage. Rest of the culture (∼980 µl) was added directly on 500 ml of liquid LB containing ampicillin. All cultures were incubated overnight (12-15h) at 32°C. Next day, plasmid extractions were performed by using NucleoBond Xtra Midi kit (Macherey-Nagel). To ensure coverage, six electroporations for LentiCRISPRv2 and three electroporations for LentiGuide-Puro were performed and mixed after plasmid extraction which yielded 500x and 800x library coverage, respectively.

### PCR amplifications from plasmid DNA

To determine sgRNA distribution in plasmid pools (LentiCRISPRv2 or LentiGuide-Puro), plasmid DNA (pDNA) were amplified by adding Illumina compatible sequences. 10 ng template DNA were mixed with 0.5 µl Phusion High-Fidelity DNA Polymerase (NEB, USA), 10 µl 5x GC Buffer (NEB, USA), 1 µl dNTP mix (10 mM each) (Thermo Fisher,USA), 2 µl Forward Stagger Mix (10 µM) and 2 µl Reverse Index Primer (10 µM) specific to each vector backbone and Nuclease-free water (NEB,USA) up to 50 µl. Thermal cycler conditions were as follows: denaturation for 30 s at 98°C, followed by (10 s at 98°C, 15 s at 63°C, 20 s at 72°C) for 16 cycles, final extension for 4 mins at 72°C. PCR amplicons were gel extracted using NucleoSpin Gel and PCR clean-up (Macherey-Nagel, Germany) kit according to manufacturer’s instructions and quantified using Nanodrop. Next generation sequencing was performed at Genewiz (USA) by using Hiseq (Illumina) with at least 10 million reads/plasmid library.

### Cell culture

MDA-MB-231, SUM159PT and SUM149PT TNBC cell lines and HEK293T cells were kind gifts from Robert Weinberg (MIT, Boston, USA). LNCaP, 22Rv1, DU145 and RWPE-1 cells were purchased from ATCC. MDA-MB-231 and HEK293T cells were cultured in DMEM (Gibco,USA) supplemented with 10% fetal bovine serum (Gibco, USA) and %1 Penicillin/Streptomycin (Gibco,USA). SUM159PT and SUM149PT cell lines were cultured in Ham’s F12 nutrient mix (Gibco, USA) supplemented with 5% FBS, 5 µg/ml insulin (Sigma-Aldrich, USA), 1 µg/ml hydrocortisone (Sigma-Aldrich, USA) and 10 mM HEPES (Thermo Fisher, USA). Immortalized human breast epithelial cells (HMLE) were a gift from Robert Weinberg (MIT, Boston, USA) [53]. HMLE cells were cultured in MEGM medium as described [54]. All prostate cell lines except RWPE-1 were cultured in RPMI 1640 (Gibco, USA) supplemented with 10% fetal bovine serum (Gibco, USA) and 1% penicillin/streptomycin (Gibco, USA). RWPE-1 cells were cultured in Keratinocyte SFM media (Gibco, USA) supplemented with 0.05 mg/mL bovine pituitary extract and 5 ng/mL human recombinant epidermal growth factor (Gibco, USA). Cells were maintained in a humidified incubator at 37°C with 5% CO_2_ level. All cell lines were tested regularly for mycoplasma infection.

### Virus production, concentration and titration

For lentiviral packaging of EPIKOL, either Fugene 6 (Roche Applied Science) or Transporter 5 transfection reagents were used. For Fugene 6 transfection, HEK293T cells were plated as 2.5 × 10^6^ cells per 10cm plate. Next day, 2500 ng EPIKOL (either in LentiCRISPRv2 or LentiGuide-Puro backbones), 2250 ng psPAX2 (Addgene 12260) and 225 ng VSV-G (Addgene 8454) plasmids were mixed in 200 μl serum-free DMEM. DNA mixture was then added into 200 μl serum-free DMEM containing 15 μl Fugene 6. For Transporter 5 transfection, 5×10^6^ HEK293T cells were seeded onto 10cm plates. The media were refreshed with low FBS media (2% FBS in DMEM) at least 2 hrs prior to transfection. For transfection, 6 ug plasmid DNA was mixed with 5.4 ug of CMV-8.2dVPR and0.6 ug of CMV-VSV-g plasmids in 0.8 mL of 150 mM NaCl solution. 36 uL of Transporter 5 was added on top of the mixture and gently mixed. In both transfection types, mixtures were incubated for 30 minutes and distributed dropwise to 10 cm plates. Next day, transfection media were replaced by 8 ml fresh media per plate. Supernatant containing viral particles was collected 48 h and 72 h post-transfection and filtered through 45-μm filters [55]. To obtain concentrated viruses, supernatants were mixed with PEG8000 (Sigma-Aldrich, USA) (dissolved in PBS as 50% (w/v)) in 10% final concentration for overnight at 4°C. Next day, supernatants were centrifuged at 2500 rpm for 20 min at 4°C and pellets were resuspended in PBS as 100x concentrated [55]. The viral aliquots were kept in -80 until usage. Viral titers were determined on the cell line of interest. Briefly, cells were seeded as 2×10^5^ cells per well of a 6-well plate and next day incubated with 10, 1, 10^−1^, 10^−2^ μl viral supernatant in the presence of 8 μg/ml protamine sulphate (Sigma-Aldrich, USA) overnight. Next day, media with viral supernatant were replaced by fresh media. 36 hours later, each well of 6 well-plate were transferred to a 10cm plate with previously determined concentrations of puromycin (Sigma-Aldrich, USA), for that cell line for 3 days. Once the cells in uninfected wells were completely eliminated with puromycin, remaining cells in other wells were compared to uninfected/unselected parental controls. The viral volume that results in 30-50% transduction efficiency was used for the downstream experiments. For EPIKOL in LentiGuide-puro backbone, LentiCas9-blast (Addgene 52962) viruses were produced in the same manner.

### CRISPR Screen

Cas9-expressing stable cell lines were generated by transducing the cells with LentiCas9-blast virus at MOI 1 for TNBC and MOI 5 for PCa cell lines. Cells were selected with blasticidin for 5 days and maintained in blasticidin-containing media for several passages prior to library infection. Negative selection screens with EPIKOL were performed as three biological replicates. Cells were transduced with EPIKOL at low MOI (0.3-0.4) with 1000x coverage for TNBC and 500x coverage for PCa in the presence of 8 μg/ml protamine sulphate. Following three days of puromycin selection, cells were collected (8×10^6^cells for TNBC, 4×10^6^ for PCa) to serve as a reference point for baseline sgRNA distribution. Remaining cells were seeded by maintaining the indicated coverage for each line and kept in culture for 15-16 population doublings. At the end of each screen, cells were collected (8×10^6^cells for TNBC, 4×10^6^ for PCa) and stored at -80°C until genomic DNA isolation.

### Genomic DNA isolation and Nested PCR

Genomic DNA (gDNA) was isolated by NucleoSpin Tissue kit (Macherey-Nagel, Germany) for TNBC cell lines and by PureLink Genomic DNA mini kit (ThermoFischer K-1820) for PCa cell lines according to manufacturer’s instructions. For PCR amplification of TNBC gDNAs (external PCR), input gDNA amount was calculated as 250x coverage of the EPIKOL library which corresponded to 13.2 µg per sample (assuming 6.6 pg DNA per cell). For each sample, 13.2 µg gDNA was divided into four PCR tubes with 3.3 µg gDNA per 100 µl reaction. In external PCR, 3.3 µg gDNA, 1 µl Phusion High-Fidelity DNA Polymerase (NEB, USA), 20 µl 5x GC Buffer (NEB, USA), 2 µl dNTP mix (10 mM each) (Thermo Fisher,USA), 5 µl Forward External Primer (10 µM), 5 µl Reverse External Primer (10 µM) and Nuclease-free water (NEB,USA) up to 100 µl were mixed on ice. Thermal cycler conditions were as follows: denaturation for 3 mins at 95°C, followed by (25 s at 95°C, 20 s at 65°C, 15 s at 72°C) for 17 cycles, final extension for 3 mins at 72°C. PCR reactions were then combined. For internal PCR, 5 µl from combined PCR products were used as a template with 5 µl Forward Stagger Mix (10 µM) and 5 µl Reverse Index Primer (10 µM) in a 100ul reaction. Thermal cycler conditions were the same as external PCR except that amplification was carried out for 23 cycles instead of 17. Final amplicons from duplicate internal PCRs were gel extracted using NucleoSpin Gel and PCR clean-up (Macherey-Nagel, Germany) kit according to manufacturer’s instructions and quantified using Nanodrop. All primer sequences are available in the **Supplementary Table 2**. Next generation sequencing was performed at Genewiz (USA) with at least 10 million reads/sample.

For library preparation of PCa samples, 4 µg of genomic DNA in total was amplified by using Kapa HiFi HotStart ReadyMix (Roche KK2602). For external PCR step, 8 reactions were prepared in 25 µL reaction volume using 0.5 µg of genomic DNA with 12.5 µL of Kapa HiFi HotStart ReadyMix, 2.5 µL of external forward primer (10uM), 2.5 µL of external reverse primer (10uM) and nuclease free water up to 25 µL. External PCR products for each sample were pooled to be used in internal PCR. For internal PCR, 2 reactions were prepared in 50 µL reaction volumes with 1 µL of pooled external PCR product, 25 µL of Kapa HiFi HotStart ReadyMix, 2.5 µL of mixed forward staggered primer pool (10uM), 2.5 µL of indexed reverse primer (10uM) and 19 µL of nuclease free water. PCR reactions were performed in the following conditions: initial denaturation at 95°C for 3 mins, denaturing at 95°C for 25 sec, annealing at 65°C for 20 sec, extension at 72°C for 15 sec. Final extension was performed at 72°C for 3 mins. The number of PCR cycles were 25 x for external PCR and 15x for internal PCR.

### Screen Analysis

MAGeCK algorithm (version 0.5.8) was used to identify significantly changed sgRNAs in knockout screens [56]. Reads from R1 fastq files were counted at sgRNA level and normalized to library size. Biological replicates were presented as individual input files during sgRNA counting. Individual counts were combined as one output count for each sgRNA in every condition with median normalization to obtain gene level log fold changes. p<0.05 cutoff was applied to gene level analysis to identify significantly depleted genes. sgRNA counts were also normalized as Read Per Million (RPM) and converted to Log_2_ values [38, 57]. Density plots of the Log_2_ transformed sgRNA counts were plotted with R using the geom_density function of the ggplot2 package to generate kernel density estimation (KDE) plots. Pearson correlations were calculated and plotted with R using the pairs.panels function in the psych package. Cumulative density plots were plotted with stat_ecdf function in ggplot2 package in R.

### Area Under the Curve Analysis

Library performance was evaluated by Area Under the Curve (AUC) calculation of predefined sgRNA groups, such as ‘Essential’, ‘Non-Essential’ and ‘Non-Targeting’ [38]. Same method was used to evaluate the performance of EPIKOL. The python code “AUC Calculation (https://github.com/mhegde)’ was downloaded from the source. AUC calculation was run following the instructions on GitHub (https://github.com/mhegde/auc-calculation). EPIKOL library specific ‘Input File’ and ‘ChiP File’ were prepared, meanwhile sgRNAs were grouped as ‘Targeting’, ‘Essential’, ‘Non-Targeting’ and used for preparation of ‘Gene Set’ files. All calculated AUC data were furtherly plotted and analyzed in GraphPad Prism 8.

### Dual-color competition assays

For validation of EPIKOL screen candidate hits, dual color competition assays were performed. Cas9-stable cells were transduced with either PGK-H2BmCherry (Addgene #21217) or PGK-H2BeGFP (Addgene #21210) viruses at high MOI ∼5 to make sure every cell was fluorescently labeled. 50,000 cells were seeded in 12 well-plates, mCherry+ cells were transduced with LentiGuide-NT1 viruses while eGFP+ cells were transduced with viruses carrying sgRNA-X for selected genes. For each gene, 2 different sgRNAs were used. After 16 hours, viral media were changed with fresh media and next day puromycin selection was started. After 3 days of puromycin selection, mCherry+ and eGFP+ cells were mixed in a 1:1 ratio and re-seeded into 24-well plates in triplicates. One day after seeding, Day0 measurements were taken by acquiring 3×3 images with a 4x objective in Cytation5 (BioTek, USA). Cells were incubated for the subsequent 16 days and images were taken at Day4, Day8, Day12 and Day16. Number of mCherry+ and eGFP+ cells were counted from images using Gen5 software (BioTek, USA) and each measurement was normalized to Day0 to determine the percentage of GFP-positive cells.

### Clonogenic Assays

To assess relative cell fitness, control cells (LentiGuide-NT1 infected) and cells carrying sgRNAs against candidate genes were seeded as 750 cells/well in triplicates in 6-well plates. Cells were allowed to grow for 12 days; media were changed regularly. At the end of the incubation period, media were discarded, cells were washed with PBS and fixed with ice-cold 100% methanol for 5 minutes. Methanol was discarded and cells were stained with crystal violet for 15 minutes. Counting of colony numbers was performed by using ImageJ with the same threshold value for each well.

### Annexin V Staining

Annexin V staining was performed with Muse® Annexin V & Dead Cell Kit (Luminex, MCH100105) according to the manufacturer’s instructions. Briefly, MDA-MB-231 cells were collected 9 days post-transduction and adjusted to be 300-500 cells/µl per gene for the measurements. Cells were centrifuged at 1200 rpm for 5 minutes. The supernatant was removed, and pellet was resuspended in 500 µl of cold PBS with 1% FBS. Cell suspension was centrifuged again and resuspended in 75 µl of cold PBS with 1% FBS and mixed with 75 µl of Annexin V & Dead Cell Reagent. Samples were incubated at room temperature for 20 minutes and analyzed with Muse Cell Analyzer (Merck, Darmstadt, Germany) with 5000 events per sample. Gates were determined according to parental cells.

### Cell Cycle Analysis

1×10^6^ MDA-MB-231 cells were collected 9 days post-transduction and centrifuged at 1200 rpm for 5 minutes. The supernatant was removed, and the pellet was resuspended in 1 ml PBS. Samples were centrifuged at 1200 rpm for 5 minutes. Most of the supernatant was removed and the pellet was resuspended in PBS. Cell suspension was added drop by drop into the freshly prepared, cold 1 ml 70% Ethanol while vortexing for fixation. Samples were kept in -20°C for 24 hours. After incubation, 200 µl of fixed cells were transferred into 15 ml conical centrifuge tube and following two rounds of wash by centrifugation at 1200 rpm for 5 minutes at room temperature, cells were resuspended in 150 µl of Muse Cell Cycle Reagent (The Muse® Cell Cycle Kit (Luminex, MCH100106)). Samples were incubated with the reagent at room temperature for 30 minutes in the dark and run through the Muse Cell Analyzer (Merck, Darmstadt, Germany) with 10,000 events per sample and analyzed with the Muse Cell Analyzer software. Gates were determined according to parental cells.

### Statistical Analysis

Analysis of EPIKOL data was performed by using RRA method in MAGeCK. Unless otherwise stated, P values were determined by two-tailed Student’s *t*-test for all experiments in GraphPad Prism8, \**P*<0.05, \*\**P*<0.01, \*\*\**P*<0.001.

### DATA availability

EPIKOL screen sequencing data are deposited to the NCBI GEO database with the accession number GSE173892.

## RESULTS

### Generation of EPIKOL and library performance in multiple cancer cell lines

To study the effect of epigenetic modifiers in multiple cancer types, we generated an epigenome-wide pooled CRISPR library. <u>Epi</u>genetic <u>K</u>nock-<u>O</u>ut <u>L</u>ibrary (EPIKOL) includes 7870 sgRNAs targeting 719 epigenetic modifiers, 25 context-specific controls and 35 pan-essential genes along with 80 non-targeting controls **(Figure 1A, B)** in two different lentiviral backbones. Both the plasmid pool and library-transduced cells were sequenced to confirm library complexity and sgRNA distribution **(Figure S1A, Figure 1C)**. sgRNA representations between the plasmid pool and transduced cell lines at the initial timepoint of the screens were highly correlated (R=0.83 for MDA-MB-231, R=0.91 for LNCaP), indicating that no bias was introduced during transduction or cloning steps **(Figure 1D, Figure S1B)**. To evaluate the efficacy of the screens, we compared the depletion scores of epigenetic-targeted genes versus essential genes and non-targeting controls. We observed significant depletion upon knockout of essential genes and no change in non-targeting controls **(Figure 1E, Figure S1C)**.

**Figure 1.**
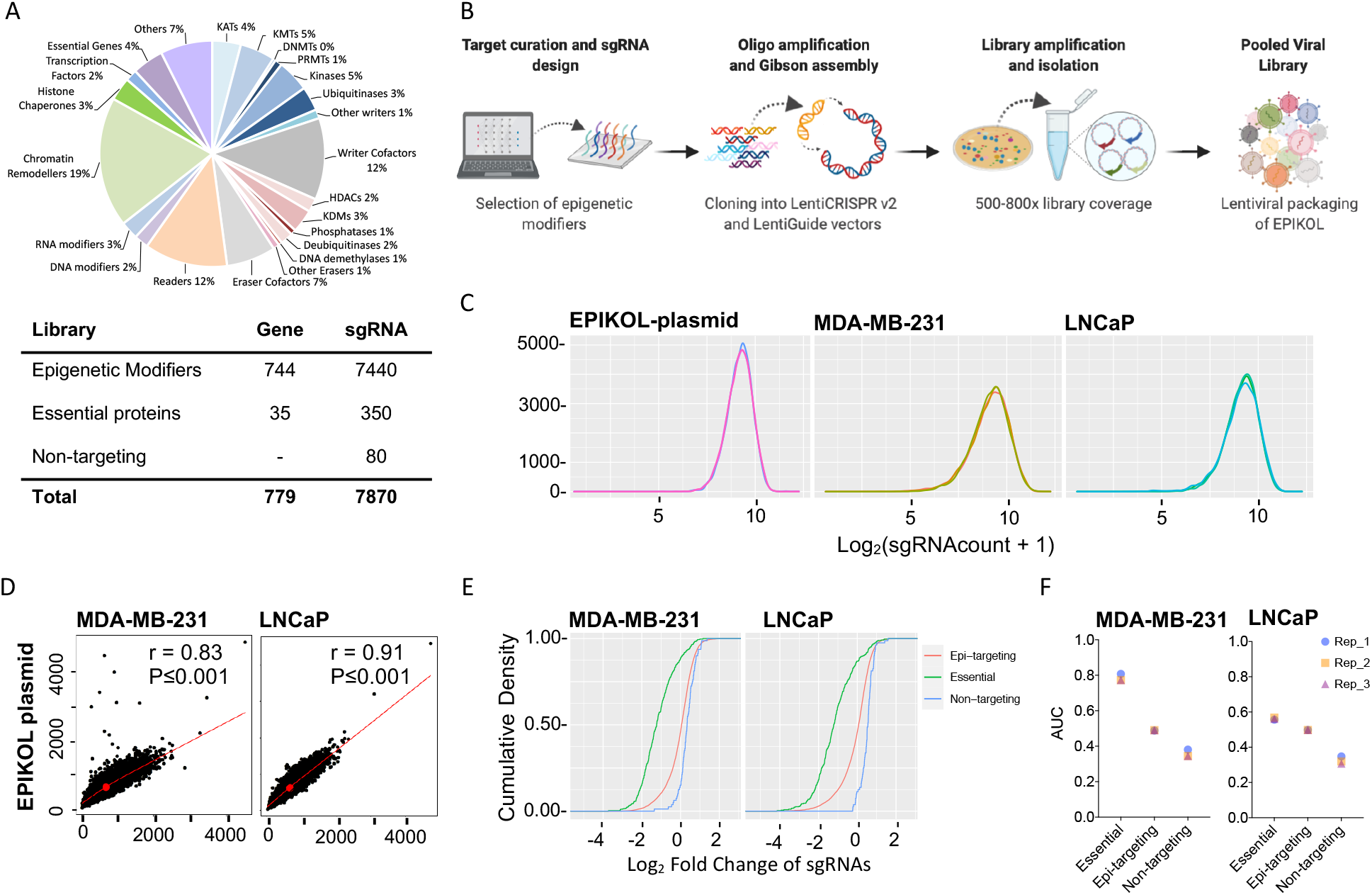
Focused Epigenetic knock-out library (EPIKOL) generation and quality check. **A.**Composition of EPIKOL library and number of sgRNAs/gene. **B**. Steps of library generation. Figure created with BioRender.com **C**. sgRNA density plots from LentiGuide plasmid containing EPIKOL and MDA-MB-231 or LNCaP cells infected with EPIKOL virus. For the cell lines, cell pellets collected after puromycin selection were used. **D**. Correlation analysis of plasmid library and samples coming from EPIKOL-infected cells at initial timepoints. **E**. Cumulative density plots showing differential depletion of sgRNAs targeting essential genes when compared to non-targeting sgRNAs. **F**. Comparison of Area Under the Curve (AUC) for sgRNAs targeting essential genes, epigenetic modifiers and sgRNAs that are non-targeting. Individual replicates of cells screened with EPIKOL for approximately 15 population doublings were shown.

Library performance was evaluated by calculating the area under the curve (AUC) for sgRNAs targeting essential genes and non-targeting controls. In multiple cell lines, essential gene targeting sgRNAs had AUC>0.5 indicating that these genes were preferentially depleted during the screens whereas non-targeting gRNAs had AUC<0.5 indicating their stationary behavior **(Figure 1F, Figure S1D)** [38]. Altogether, these initial quality check analyses demonstrated that EPIKOL preserves normal distribution of sgRNAs both in plasmids and infected cells and functions as expected in depletion screens.

### EPIKOL screens revealed epigenetic vulnerabilities of TNBC and prostate cancer cell lines

To uncover epigenetic modifiers important for cancer cell fitness, we conducted negative selection (drop-out) screens using EPIKOL. Three different triple negative breast cancer (TNBC) cell lines MDA-MB-231, SUM149PT and SUM159PT were screened in addition to non-malignant human mammary epithelium cells (HMLE) [53]. Similarly, prostate cancer (PCa) cell lines LNCaP, DU145 and 22Rv1 were screened along with the normal-like immortalized prostate epithelium cell line RWPE-1. In each screen, Cas9-expressing cell lines were transduced with EPIKOL in the lentiGuide vector at a low multiplicity of infection (MOI 0.3-0.4) with at least 500x coverage to ensure that every cell carries one sgRNA and each sgRNA is represented in at least 500 cells to maintain the complexity throughout the screen **(Figure 2A)**. Following puromycin selection, transduced cells were cultured for 15-16 population doublings by maintaining the sgRNA coverage at each passage. PCR amplified sgRNA barcodes from the initial and final timepoints of the screen were analyzed by deep sequencing. To determine relative sgRNA abundance at each timepoint, raw read counts were normalized to reads per million and log2 transformed **(Figure 2B, Figure S2A)**. Model-based Analysis of Genome-wide CRISPR/Cas9 Knockout (MAGeCK) was performed to determine gene-level depletion scores using median normalization and determine the epigenetic modifiers that decrease cell fitness. A number of epigenetic modifiers were found to be significantly depleted in TNBC cell lines MDA-MB-231 (140), SUM149PT (140) and SUM159PT (98). Similar numbers of epigenetic modifiers were also depleted in PCa cell lines LNCaP (148), DU145 (181) and 22Rv1 (173) **(Figure 2C, Figure S2B)**. Among these, epigenetic modifiers that were previously implicated in breast cancer cell fitness such as PRMT5 [58, 59], HDAC3 [60], NPM1 [61, 62] were depleted in MDA-MB-231 cells serving as positive controls. Similarly, for prostate cancer, KDM1A [63], BRD4 [64], and PRMT1 [65] were depleted in LNCaP cells as well as AR, FOXA1 and NCOA1 [66, 67], thus serving as positive controls. In addition, well-known cancer survival genes such as PELP1 and PRMT family members were identified as common hits in all the six cancer cell lines screened by EPIKOL **(Figure 2D)** [58, 59, 68-70]. These results indicated that our epigenetic-focused screening approach is able to identify genes critical for cancer cell viability. Therefore, we first focused on characterizing novel hits from the TNBC screen, which have not been previously linked to TNBC cell viability.

**Figure 2.**
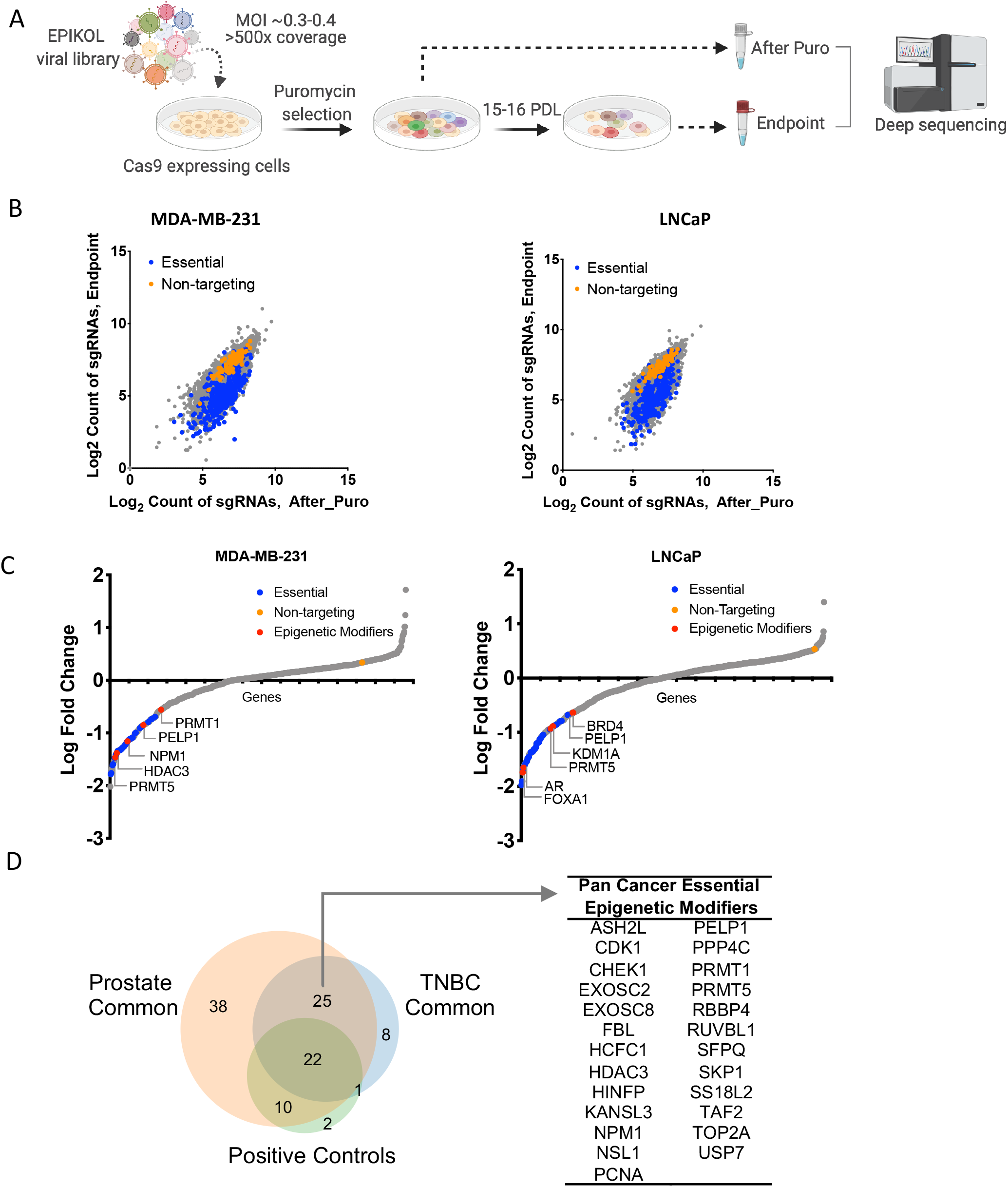
EPIKOL screens on TNBC and prostate cancer cell lines revealed cancer-specific and pan-cancer epigenetic modifiers that regulate cell fitness. **A**. Summary of screening procedure. Figure created with BioRender.com **B**. Log2 counts of sgRNAs at initial and final time points **C**. Log fold changes of genes after screening with EPIKOL for at least 15 population doublings. **D**. Common hits of EPIKOL screens on TNBC (MDA-MB-231, SUM159PT, SUM149PT) and Prostate cancer cell lines (LNCaP, DU145, 22Rv1) identified in p<0.05 cutoff.

### Effects of novel candidate genes on TNBC cell fitness were validated in dual-color competition assay

In order to validate the results of EPIKOL screens, we first identified the genes that were commonly depleted in at least two TNBC cell lines but not significantly depleted in the control HMLE cells **(Figure 3A)**. From this, 15 genes were found to be essential in all TNBC cell lines including some of the well-known regulators of cancer cell fitness such as UHRF1 [71], PELP1 [72] and PRMT1 [73]. In total, 40 genes (including several controls) were selected for further *in vitro* validation experiments based on their depletion p-values, log fold changes and gene rankings in different screens. 2 sgRNAs per gene were cloned individually into lentiGuide-puro vector and a dual-color competition assay was performed in all TNBC cell lines **(Figure 3B)**. In this assay, GFP-positive cells were transduced with sgRNAs targeting a hit gene and admixed in a 1:1 ratio with mCherry-positive cells transduced with a non-targeting sgRNA. The admixed cell populations were then monitored using a fluorescence cell imaging platform for 16 days to assess relative cell fitness. Cells carrying sgRNAs targeting a hit gene (eGFP+ cells) were outcompeted by the cells carrying non-targeting (NT1) control gRNA (mCherry+ cells) in almost all cell lines tested **(Figure S3)**. Of note, depletion ratios varied depending on the cell type; the most significant depletion was observed on MDA-MB-231 followed by SUM149PT and SUM159PT which was in line with the depletion ratios observed during EPIKOL screens. The competition assay indicated that shared members of MLL/COMPASS complexes (ASH2L, WDR5, RBBP5) as well as the NuA4 (YEATS4, VPS72) and NSL complex members (KANSL2, KANSL3, KAT8) have strong effects on the fitness of TNBC cell lines. Collectively, these findings show that EPIKOL screens identified novel epigenetic modifiers that regulate triple-negative breast cancer cell fitness.

**Figure 3.**
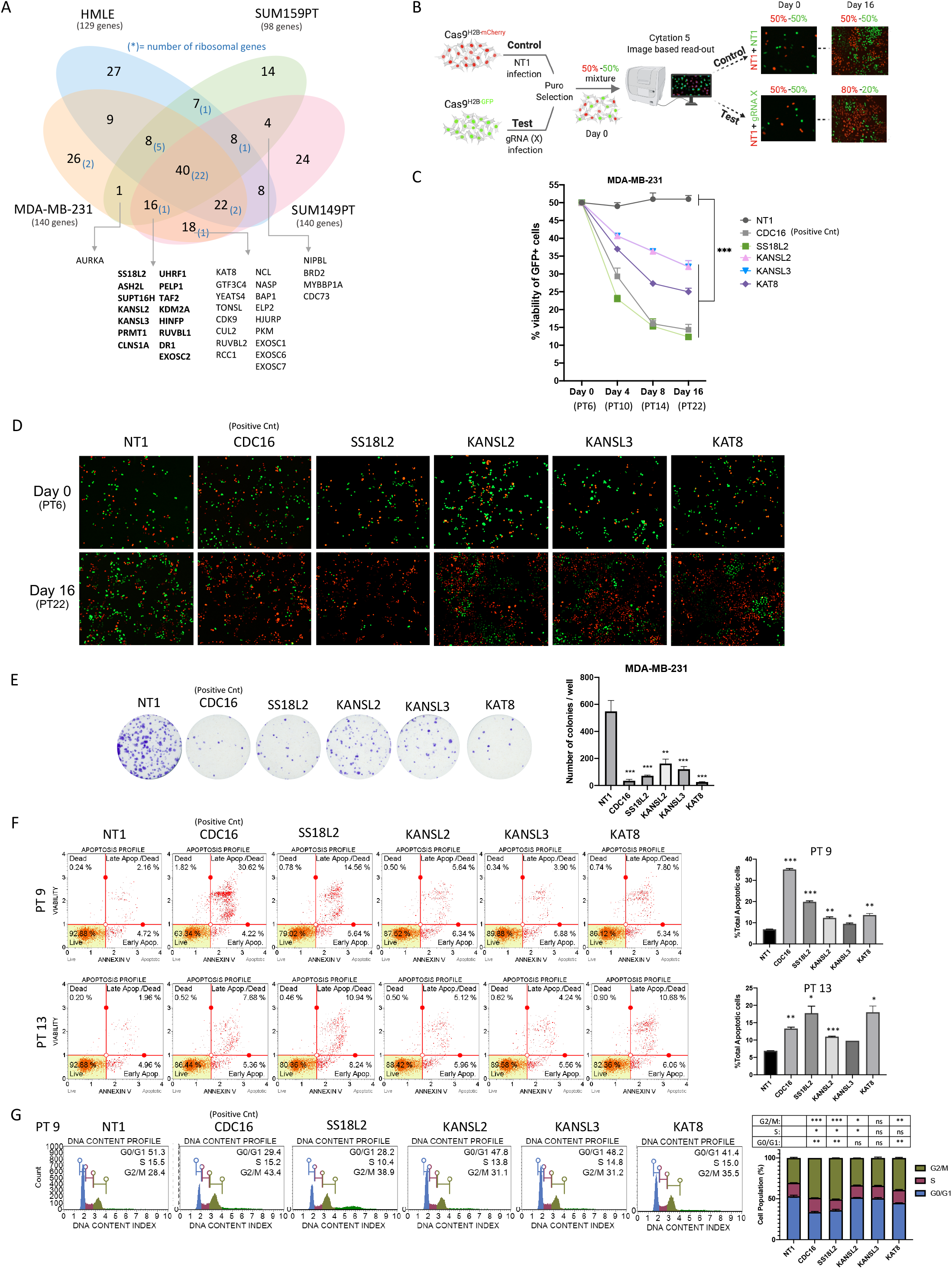
Effects of candidate genes on MDA-MB-231 fitness were validated with functional assays *in vitro*. **A**. Venn diagram showing cell line specific or common genes that are found in p<0.05 cutoff. 15 genes in bold show TNBC specific epigenetic modifiers that were depleted in all three TNBC cell lines. Others are the genes that were commonly depleted in 2 different TNBC cell lines but not in HMLE **B**. Summary of dual-color competition assay for *in vitro* validation of candidate epigenetic modifiers. **C**. Results of dual-color competition assay for selected hits in MDA-MB-231 cells. PT: post-transduction day. **D**. Representative images taken with Cytation5 at Day0 and Day16 of competition assay for MDA-MB-231 cells. mCherry+ cells were infected with Non-targeting sgRNA (NT1) as control while eGFP+ cells were infected with sgRNA targeting the gene of interest. **E**. Representative images of long-term clonogenic assay for MDA-MB-231 cells infected with sgRNAs against selected hits. **F**. Annexin V & dead cell assay results of selected genes on two different time points and their statistical analysis. **G**. Cell cycle analysis of selected genes on PT9 and its statistical analysis. P values determined by two-tailed Student’s *t*-test in comparison to NT1; *P<0.05, **P<0.01, ***P<0.001

### Knockout of individual epigenetic modifiers caused growth defects in TNBC cell line MDA-MB-231

To further delineate the effects of novel epigenetic modifiers that regulate cell fitness, four of the TNBC specific genes (SS18L2, KANSL2, KANSL3 and KAT8) were selected based on their strong depletion scores in MDA-MB-231. Three of these genes belong to the same complex, namely the *non-specific lethal* (NSL) complex. KANSL2 and KANSL3 are structural components of NSL complex together with KANSL1. KAT8 (*MOF, MYST1*) is the catalytic member of the complex and acetylates histone lysine residues [74]. On the other hand, SS18L2 is the homolog of the *SS18* gene, which is associated with chromosomal translocation characteristics of synovial sarcoma. However, the exact role of SS18L2 in synovial sarcoma or any other cancer is not known [75]. NSL complex members (KANSL2, KANSL3, KAT8) and SS18L2 showed a strong TNBC-specific effect in EPIKOL screens. In competition assays, cells carrying sgRNAs targeting all four hit genes were significantly depleted in MDA-MB-231 cells over 16 days **(Figure 3C, D)**. Long-term colony formation assays clearly showed that knockouts of all selected genes exert strong fitness defects when compared to the control sgRNA-expressing cells **(Figure 3E)**. Suppression of these genes led to 65-75% fewer number of colonies compared to control conditions with the SS18L2 and KAT8 depletion phenotype reaching to the level observed with the depletion of positive control CDC16. Taken together, these results show that knocking out either SS18L2 or members of NSL complex have a profound effect on TNBC cell fitness.

### Knockout of epigenetic modifiers induced apoptosis in MDA-MB-231 cells

To identify the mechanism through which cell fitness is reduced, we first investigated whether knockout of these genes result in apoptosis. Annexin V & Dead cell staining showed significantly more cells in early- and late-apoptotic states upon knockout when compared to control cells **(Figure 3F)** at two different timepoints. On post-transduction (PT) day 9, knockout of SS18L2 induced apoptosis significantly in line with the effect that we observed in the first four days of competition assays **(Figure 3D)**. While the knockout of NSL complex members induced apoptosis significantly at PT9, their effects, especially of KAT8, were more pronounced at PT13.

We also observed a reduced number of cells in the G0/G1 and S phases of cell cycle and accumulation at G2/M phase upon SS18L2 and KAT8 knockout **(Figure 3H)**. This indicates that knockout of these genes may result in mitotic arrest. Collectively, these findings suggest that four candidate genes are essential to TNBC cells, which might be exploited for therapeutic purposes. These proof-of-principle experiments demonstrate that our focused epigenome-wide CRISPR library, EPIKOL, is an easy-to-use functional genomics tool that enables identification of epigenetic modifiers important for cancer cell fitness.

## DISCUSSION

In this study, we present a focused epigenetic knockout library (EPIKOL) that can be utilized to investigate chromatin-based vulnerabilities in different biological contexts. We performed eight screens in 2 cancer types and identified novel chromatin modifiers that regulate prostate and triple-negative breast cancer cell fitness. Successful validation of individual candidate genes through follow-up competition and cell growth assays showed that EPIKOL is suitable to functionally interrogate a wide range of epigenetic modifiers. In contrast to most currently available epigenome-focused libraries, which only target chromatin modifiers such as writers, readers and erasers [44-46], EPIKOL targets a wider range of genes coding for chromatin complex cofactors and structural components [48]. Thus, its use will likely lead to a broader understanding of the functions of these complexes as a whole.

EPIKOL is available both in LentiCRISPRv2 and LentiGuide backbones, both of which showed normal distribution of sgRNAs, indicating that all elements of the library are present equally in both backbones in pooled format. Availability of EPIKOL in LentiCRISPR v2 backbone might expedite the screening process by eliminating the need for prior Cas9 introduction especially in patient-derived xenograft models and primary cell lines, in which the culturing time of the material is limited. In such cases, smaller library size will also reduce the amount of initial material required to maintain the complexity.

Library performance as evaluated by AUC calculations for non-targeting sgRNAs and sgRNAs targeting essential genes were in line with the literature [38], suggesting that EPIKOL is an efficient loss-of-function library with highly effective controls. Another advantage of EPIKOL is the presence of sgRNAs targeting context-specific control genes from different families such as nuclear receptors, transporters and EMT-related proteins. Depletion of sgRNAs targeting such genes provides another layer of confidence for the quality control of the screens depending on the investigated phenotype. For example, Androgen Receptor (AR) targeting sgRNAs were significantly depleted in AR-dependent prostate cancer cell lines LNCaP and 22Rv1 while no change was observed in AR-negative cell line DU145 and TNBC cell lines suggesting that EPIKOL can distinguish tissue or cell line specific hits.

From drop-out screens in multiple cell lines, we identified novel epigenetic modifiers for cancer cell fitness as well as the previously studied ones such as PRMT5, HDAC3, FOXA1 and LSD1 [58-60, 67]. The comparison between six different cancer cell lines revealed 25 epigenetic modifiers commonly depleted in all cell lines tested. Among them, several genes belong to the same complex such as PRMT family, exosome complex and MLL complexes highlighting the role of these epigenetic complexes as pan-cancer essential epigenetic modifiers. Interestingly, some of these genes were previously identified as common essential genes in DepMap based on their significant depletion in almost 750 different cancer cell lines proving that EPIKOL screens can successfully identify common epigenetic pathways in cancers in addition to cell line or cancer specific ones [76]. We also identified ASH2L as a common essential gene in both cancer types through EPIKOL screens, while it was not classified as a common essential gene in previously performed screens.

SS18L2 was one of the genes that drew our attention since it was significantly depleted in all TNBC EPIKOL screens. Competition assays confirmed the strong effect of SS18L2 knockout on TNBC cell survival. SS18L2 is a homolog of SS18 gene which is associated with the malignant gene translocation in synovial sarcoma [75]. The role of SS18L2 is not known in cancers and there is no study showing the effect of this gene in TNBC. Here, for the first time, we showed that knockout of SS18L2 decreased TNBC cell survival, induced G2/M arrest and concomitantly induced apoptosis. In the future, it will be important to study how SS18L2 exerts this strong effect on TNBC cells and whether it can serve as a target during treatment of this cancer type.

Another group of epigenetic modifiers that we identified as important for TNBC cell fitness belong to the NSL complex. Core members KANSL2 and KANSL3 were specifically depleted in EPIKOL screens of 3 different TNBC cell lines while they did not show significant effects in HMLE cells. Importantly, KAT8, the catalytic subunit of the NSL complex, was also a common hit in MDA-MB-231 and SUM149PT cell lines. We showed that TNBC cells were dependent on NSL complex through several functional assays. NSL complex mainly regulates H4K16Ac which is necessary for chromatin relaxation and is present at active enhancers and promoters [77, 78]. Downregulation of KAT8 and reduced H4K16Ac are associated with different cancer types, such as ovarian cancer [79], colorectal carcinoma [80], hepatocellular carcinoma [81] and gastric cancer [80]. In contrast, increased expression of KAT8 in non-small cell lung cancer results in poor survival [82]. Analyses of 298 cases of primary breast carcinomas through immunohistochemistry indicated that KAT8 protein level was decreased only in small subset of the cases (18%) indicating that the role of KAT8 might be different in breast cancer development than in other types of cancers [83]. To the best of our knowledge, there is no study showing how KAT8 and NSL complex regulate TNBC cell fitness and survival. Here, as a result of EPIKOL screens, we showed that KAT8 together with the two structural components of NSL complex (KANSL2 and KANSL3), decrease TNBC cell fitness and induce apoptosis. Since KATs can also acetylate non-histone proteins, it will be important to dissect out the mechanisms by which KAT8 regulates TNBC cell fitness to be able to precisely target this complex in breast cancer. Interestingly, previously published epigenome-wide libraries did not include KANSL2, KANSL3 and SS18L2 targeting sgRNAs, and therefore failed to identify strong effects of these genes on cell survival [44-46]. Therefore, the broad targeted gene list of EPIKOL ensures identification of true-positive hits.

Altogether, we generated and validated a focused epigenetic CRISPR library that offers a powerful platform to identify critical epigenetic modifiers. Epigenetic modifying enzymes are promising therapeutic targets to treat cancer as they regulate numerous critical cellular responses including cell growth, metastasis, apoptosis and others. The smaller library size both allows for CRISPR screens in various physiological relevant models, where cell numbers are extremely limited, and also provides a focused perspective in terms of identifying epigenetic complexes that might be responsible for the development of aggressive phenotype in cancer. EPIKOL is a robust functional genomics platform to interrogate chromatin modifiers and can guide the discovery of cell-type specific epigenetic vulnerabilities of cancers.

## Author Contributions

Study design: TB-O, OY-B, BG, TTO and NAL; data generation: OY-B, BG, EYK, GK and RG; library generation: AK, TM, FU, TB-O, TTO and NAL; data analysis: ACA, ADC and TM; data interpretation: OY-B, BG, TB-O, TTO and NAL; initial manuscript draft: OYB, BG, TTO and TB-O; approved final manuscript: all authors.

## Acknowledgements

Financial support was obtained from The Scientific and Technological Research Council of Turkey (TUBITAK) (1003-216S461 Grant). The authors gratefully acknowledge the use of the services and facilities of the Koç University Research Center for Translational Medicine (KUTTAM), funded by the Presidency of Turkey, Presidency of Strategy and Budget.

## FIGURES

**Supplementary Figure 1.**
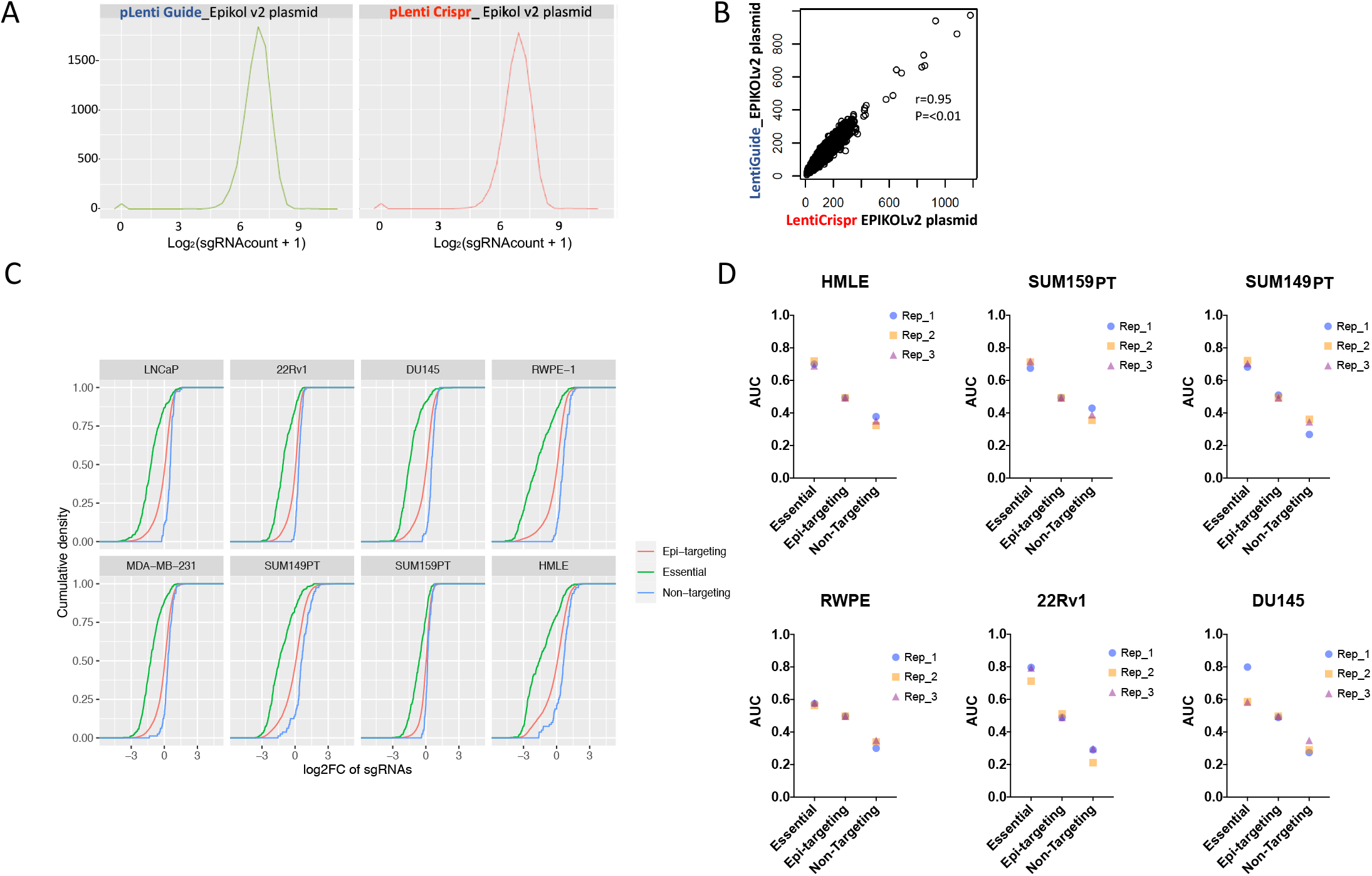
Quality Check of EPIKOL in plasmid and transduced cell level. **A**. Density and correlation plots for EPIKOL in LentiGuide or LentiCRISPRv2 backbone. **B**. Correlation analysis of EPIKOL amplified from two different plasmids. **C**. Cumulative density plots showing differential depletion of essential genes during screen when compared to epi-targeting genes or non-targeting controls **D**. Area Under the Curve (AUC) calculations for EPIKOL screens in all cell lines.

**Supplementary Figure 2.**
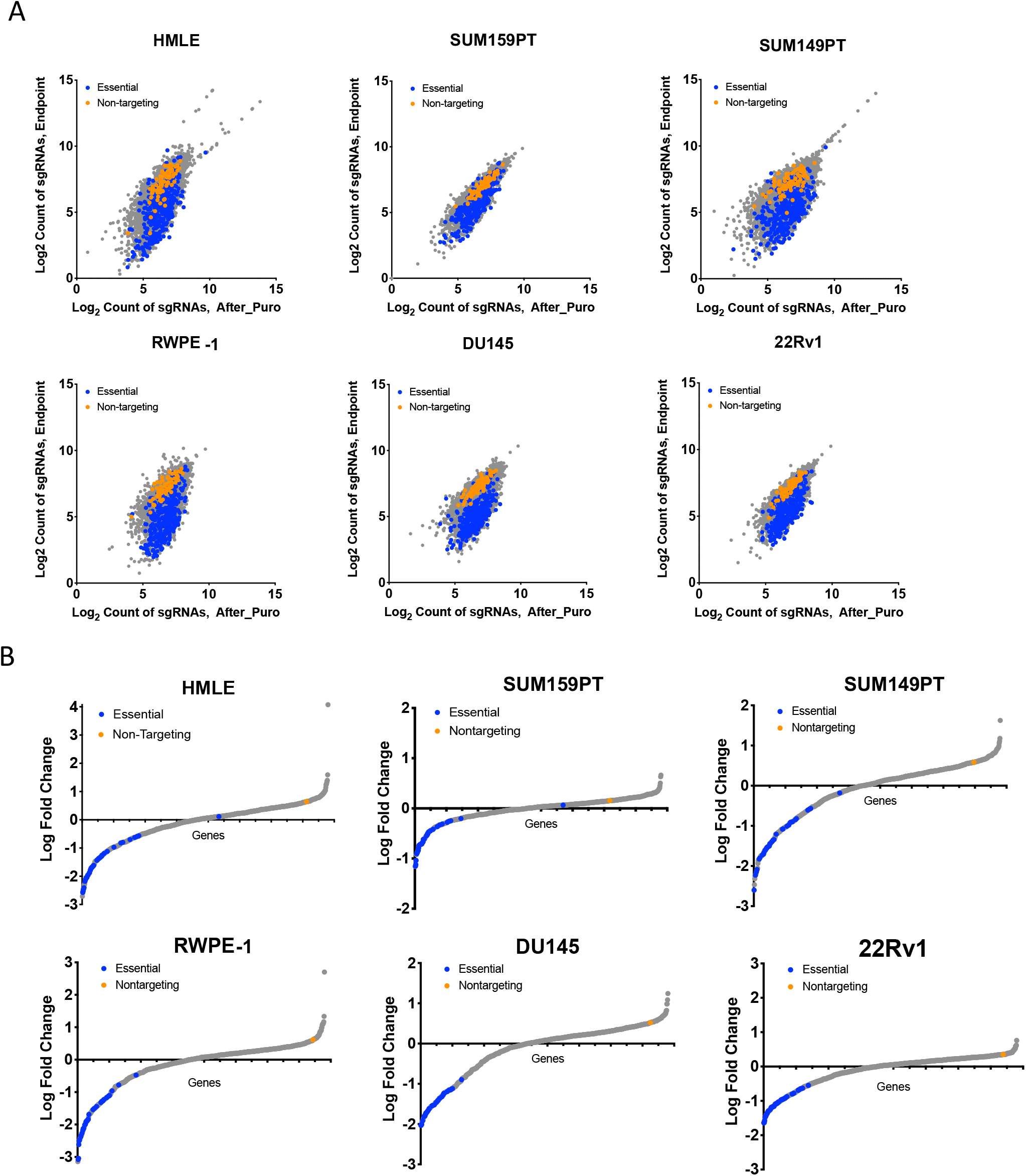
sgRNA level log2 count and gene-level waterfall plots for EPIKOL screens on each cell line. **A**. Log2 counts of sgRNAs at initial and final time points in TNBC and prostate cells lines. **B**. Waterfall plots for Log_2_ fold changes of genes after screening with EPIKOL for at least 15 population doublings in TNBC and prostate cell lines.

**Supplementary Figure 3.**
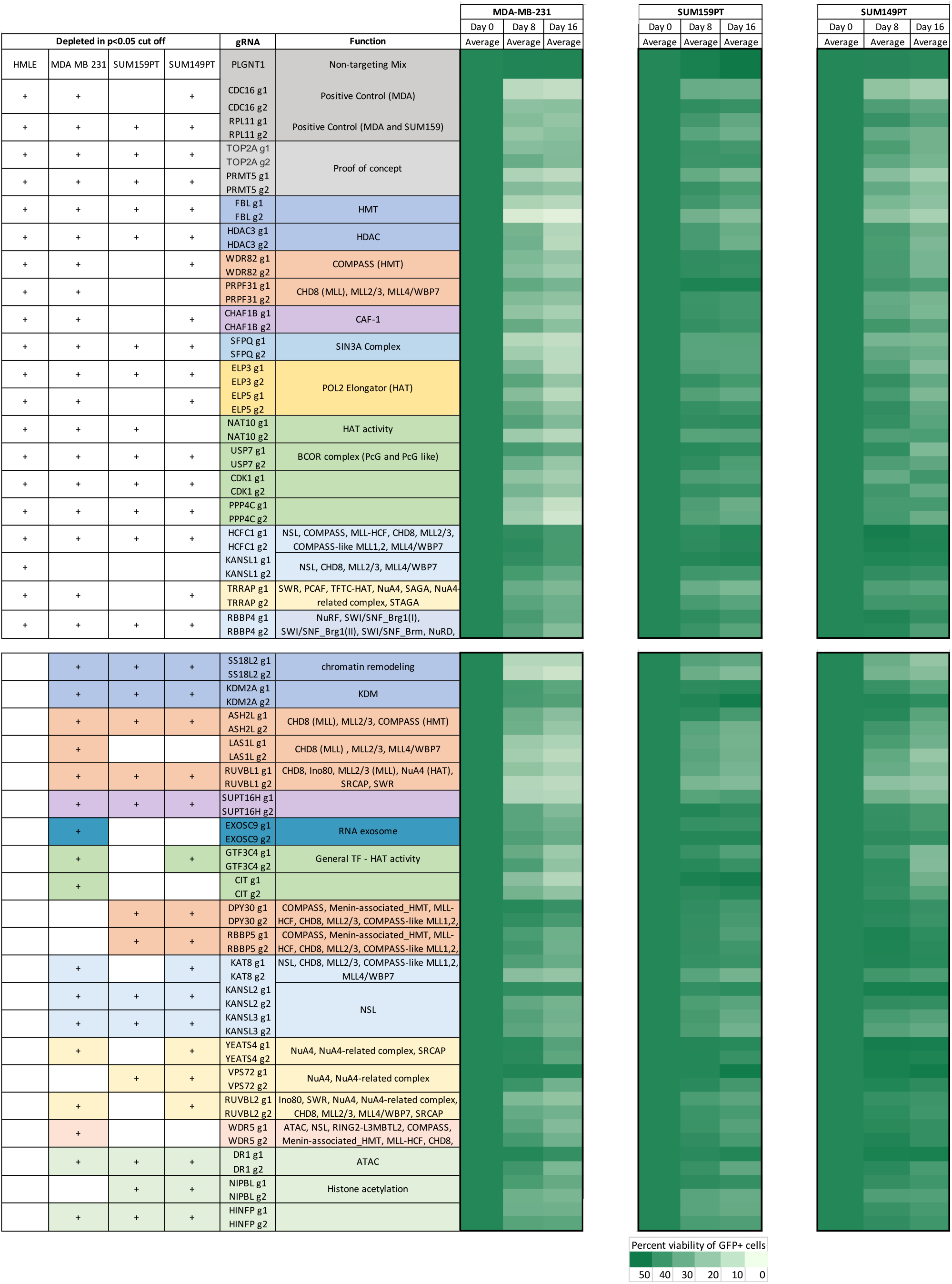
Heat-map of competition assay results normalized to Day0 for TNBC cell lines. Left-hand side shows if a given gene is found in p<0.05 cutoff in given cell line. Complexes of the genes were indicated if applicable. Day8 and Day16 measurements of all sgRNAs were normalized to the corresponding Day0 measurement.

